# Breaching the cell-envelope barriers of gram-positive and fungal microbes by a type VI secretion system in *Acidovorax citrulli*

**DOI:** 10.1101/2021.05.31.446370

**Authors:** Tong-Tong Pei, Yumin Kan, Zeng-Hang Wang, Ming-Xuan Tang, Hao Li, Shuangquan Yan, Yang Cui, Hao-Yu Zheng, Han Luo, Tao G. Dong

**Affiliations:** State Key Laboratory of Microbial Metabolism, Joint International Research Laboratory of Metabolic & Developmental Sciences, School of Life Sciences and Biotechnology, Shanghai Jiao Tong University, Shanghai, 200240, China; Department of Ecosystem and Public Health, University of Calgary, 3330 Hospital Dr. NW, Calgary, AB, T2N4Z6, Canada

**Keywords:** protein secretion, interspecies interaction, fungi, Mycobacteria

## Abstract

The type VI secretion system (T6SS) is a double-tubular toxin-injection nanomachine widely found in gram-negative human and plant pathogens. The current model depicts that the T6SS spear-like Hcp tube is powered by the contraction of an outer sheath to drill through the envelope of a neighboring cell, achieving cytosol to cytosol delivery. However, gram-positive bacteria seem to be impenetrable to such T6SS action. Here we report that a plant pathogen *Acidovorax citrulli* (AC) deploys a highly potent T6SS to kill a range of bacteria including *Escherichia coli, Pseudomonas aeruginosa*, *Bacillus subtilis*, and *Mycobacterium smegmatis* as well as fungal species including *Candida albicans* and *Pichia pastoris*. Using bioinformatic and biochemical assays, we identified a group of T6SS effectors and characterized one effector RhsB that is critical for interspecies interaction. We report that RhsB contains a conserved YD-repeat domain and a C-terminal nuclease domain. Toxicity of RhsB was neutralized by its downstream immunity proteins through direct interaction. RhsB was cleaved at the C-terminal end and a catalytic mutation within the internal aspartic protease abolished such cleavage. Collectively, the T6SS of AC displays potent activities to penetrate the cell envelope barriers of gram-positive and fungal species, highlighting the greatly expanded capabilities of T6SS in modulating microbiome compositions in complex environments.

## Main text

The COVID-19 pandemic has cost tremendous loss in human lives and global economy, revealing how fragile our society remains to infectious agents. In fact, microbial infectious diseases pose perpetual threats to public health, including tuberculosis affecting billions of people worldwide and emerging fungal pathogens causing millions of nosocomial infections (1, 2). One solution to combat infections is to learn from natural weapons that microbes use to outcompete others. One such lethal weapon is the type VI secretion system (T6SS) that gram-negative bacteria commonly employ to deliver toxins into neighboring cells through a phage-tail-like double tubular contractile structure (3–5). A spear-like tube with toxic effectors is ejected outward in milliseconds with enough power to drill through the cell wall and two cellular membranes of gram-negative cells (5). T6SS organisms show great variability in the arsenal of toxic effectors and thus killing capability against susceptible amoeba, yeast and other eukaryotic cells (3, 6–8). However, gram-positive bacteria seem to be resistant to T6SS killing.

*Acidovorax citrulli* (AC) is a gram-negative seed-borne plant pathogen that causes bacterial fruit blotch of cucurbit crops including melons and watermelons and is extremely difficult to eradicate (9). Similar to human pathogens, *A. citrulli* has rapidly spread globally in the last few decades and exhibits genetically and phenotypically distinct variants (9). Here, we report that the AC type strain AAC00-1 could employ its constitutively active T6SS to effectively outcompete a variety of bacteria and fungi including *E. coli*, *Enterobacter cloacae*, *Pseudomonas aeruginosa*, *Bacillus subtilis*, *Mycobacterium smegmatis*, *Pichia pastoris*, *Saccharomyces cerevisiae* and *Candida albicans* (Figure 1, A&B). Survival of these prey species was significantly reduced when they were exposed to wild type AC compared to the T6SS null Δ*tssM* mutant. Collectively, these data of interspecies interactions underscore the broad ecological impact of the T6SS on diverse natural microbial communities, beyond the previously known susceptible gram-negative and fungal microbes.

**Fig. 1.**
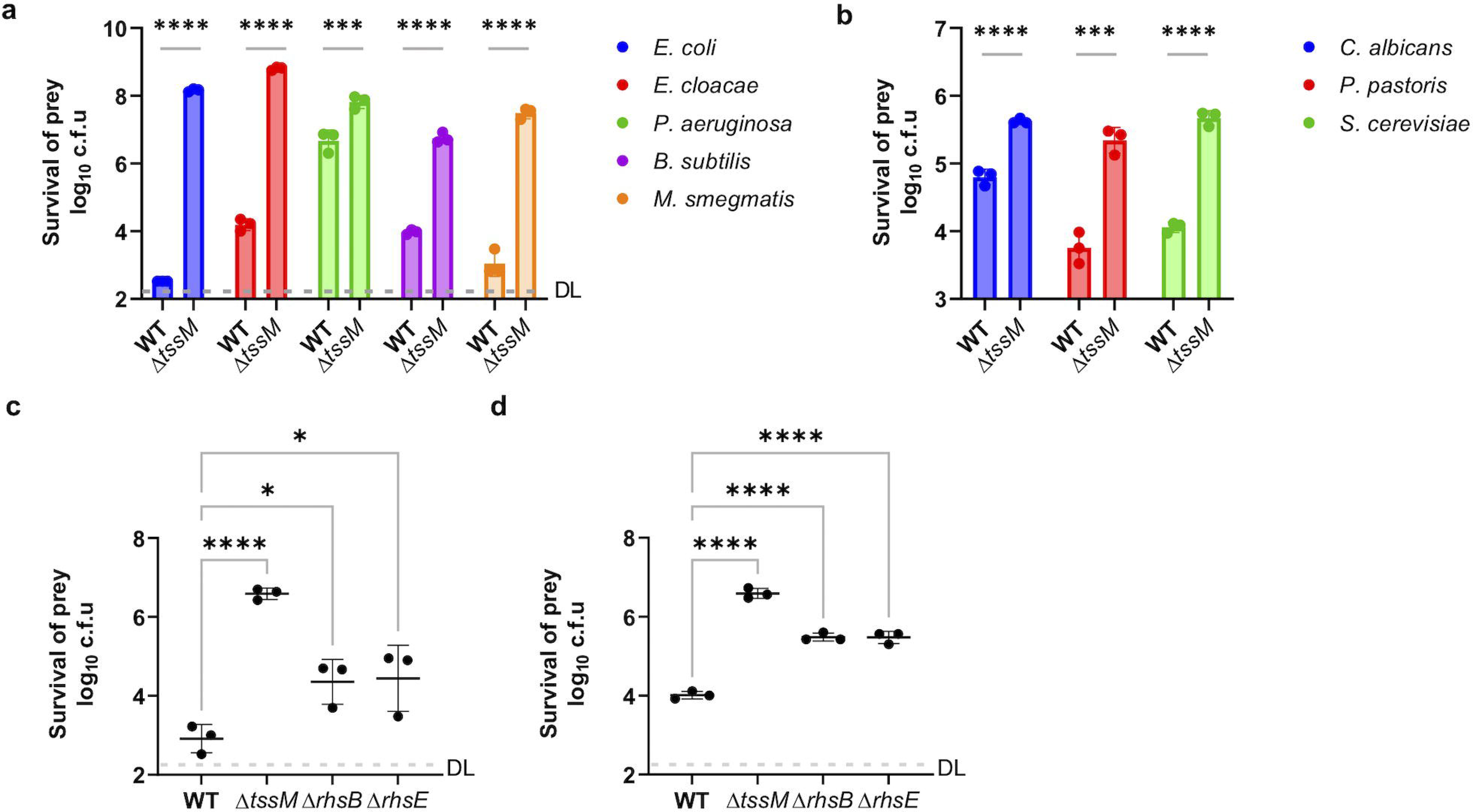
The T6SS of *A. citrulli* displays anti-bacterial and anti-fungal activities. Competition assay of wild type (WT) and the T6SS-null Δ*tssM* mutant against a panel of bacterial (A) and fungal (B) strains. Survival of *E. coli* MG1655 (C) and *B. subtilis* PY79 (D) attacked by AAC00-1 wild-type strain, Δ*tssM*, Δ*rhsB,* and Δ*rhsE* mutants for 1 h and 3 h, respectively. Survival of prey cells was determined by serial dilutions on selective media. Error bars indicate the mean +/− standard deviation of three biological replicates and statistical significance was calculated using One-way ANOVA test for each group, \**P* < 0.05, \*\*\**P* < 0.001, \*\*\*\**P* < 0.0001. DL, detection limit.

Next we decided to identify T6SS effectors responsible for the killing activities. The AAC00-1 genome contains 12 *vgrG* operons that are expected to encode at least one effector per operon (10, 11). Using bioinformatic and secretome analyses, we identified 17 VgrG-related effector genes and constructed corresponding deletion or inactivating-insertion mutants. Using bacterial competition assays against *E. coli* and *B. subtilis* prey, we screened these mutants and found two mutants (Aave_0499, named RhsB and Aave_2838, named RhsE) with impaired but not abolished killing activities (Figure 1 C&D). These results highlight a functional redundancy of effectors that collectively contribute to killing activities.

For the two mutants, both genes encode Rhs-family proteins with an N-terminal PAAR domain, a middle YD-repeat/Rhs domain, and a C-terminal domain of unknown function (Figure 2A). Using Phyre2 sequence analysis, we found no significant hit for Aave_2838 but the C-terminal domain of Aave_0499 is distantly related to a virus-type replication-repair nuclease (PDB: 4qbn) with 24% identity (12). Downstream of *rhsB* reside two small predicted genes of unknown functions sharing 69% identity and equal length in protein sequence. We name the two downstream genes *rimB1* and *rimB2* (<u>R</u>hs-<u>im</u>munity <u>B</u>), respectively.

**Fig. 2.**
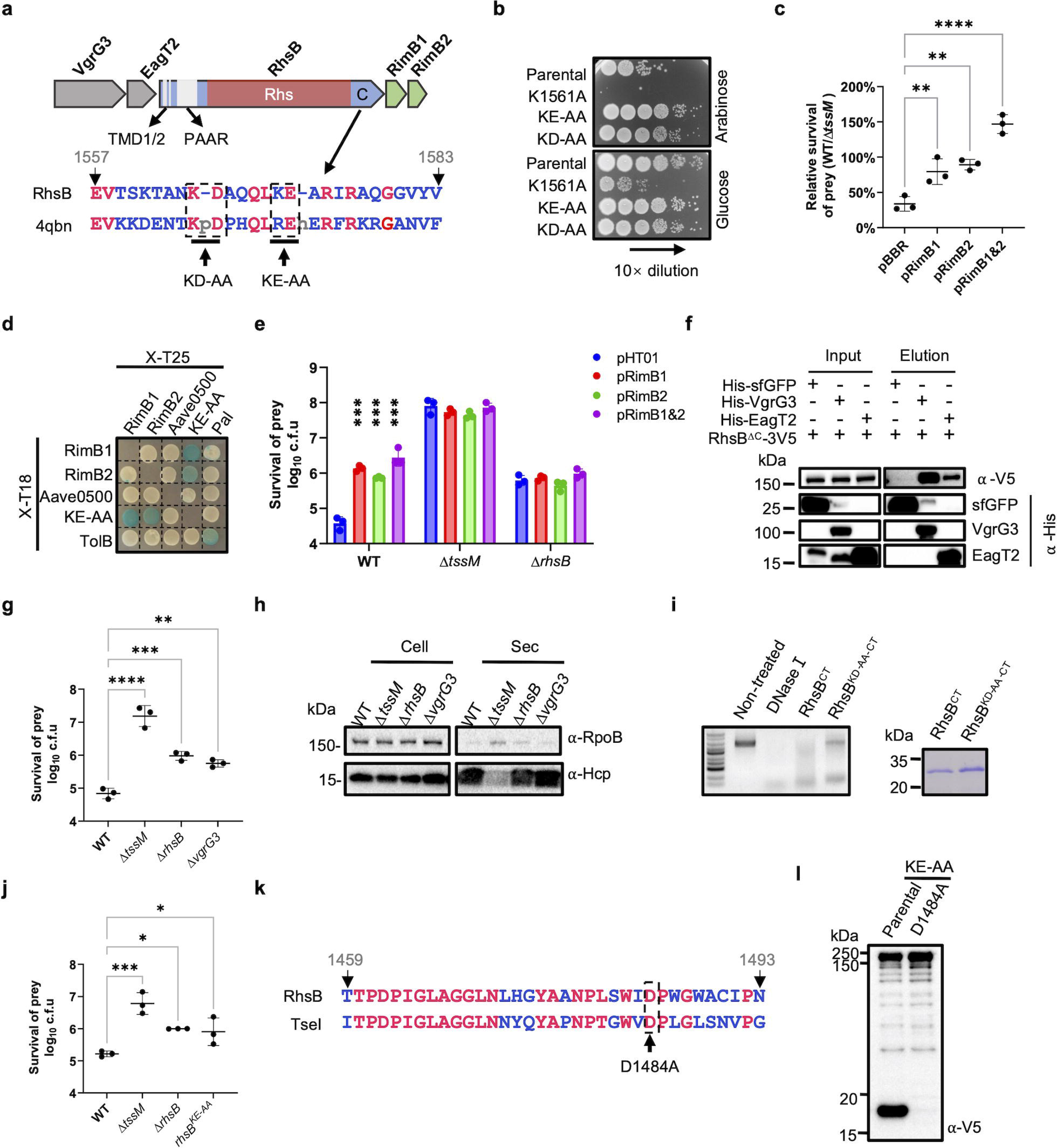
Characterization of RhsB function. (A) Operon structure of *rhsB.* The sequence of the RhsB C-terminus is aligned with the sequence of a virus-type replication-repair nuclease domain (VRR-Nuc, PDB: 4qbn). (B) Toxicity of expressing RhsB C-terminus (parental) and its catalytically inactive mutants in *E. coli*. All constructs were cloned on pBAD vectors and survival of *E. coli* was tested by serial plating on arabinose (induction) and glucose (repression) plates. (C) Competition assay of wild type (WT) and the T6SS-null Δ*tssM* mutant against the effector–immunity deletion mutant Δ*rhsB*-*rimB1/2* complemented with an empty vector (pBBR) or a vector carrying the immunity gene *rimB1*, *rimB2* or *rimB1* and *rimB2* together as indicated. The data point indicates the relative survival of prey cells attacked by wild type compared with that by T6SS mutant. Error bars indicate the mean +/− standard deviation of three biological replicates and statistical significance was calculated using a two-tailed Student’s *t*-test, \*\**P* < 0.01, \*\*\*\**P* < 0.0001. (D) Bacterial two-hybrid analysis of RhsB–RimB1/2 interaction. Proteins fused with the adenylate cyclase T25 or T18 subunits were co-expressed in the reporter strain BTH101 as indicated. (E) Competition assay of wild type (WT), Δ*tssM* and Δ*rhsB* mutant against the *B. subtilis* PY79 carrying an empty vector (pHT01) or a vector carrying the immunity gene *rimB1*, *rimB2* or *rimB1* and *rimB2* together as indicated. Cells of killer and prey were mixed at a ratio of 20:1 (killer: prey) and spotted on LB agar plates with 1mM IPTG for 3 h at 37 °C. Error bars indicate the mean +/− standard deviation of three biological replicates and statistical significance was calculated using a two-tailed Student’s *t*-test, \*\*\**P* < 0.001. (F) Interaction of RhsB ΔC with VgrG3 and chaperone EagT2. Pulldown analysis was performed using His-tagged sfGFP (control), VgrG3 or EagT2 and V5-tagged RhsB ΔC. (G) Competition assay of wild type (WT), Δ*tssM*, Δ*rhsB* and Δ*vgrG3* mutant against the *B. subtilis* PY79. (H) Secretion analysis of Hcp. Secretion of Hcp was detected by Western blotting analysis. RpoB serves as an equal loading and autolysis control. (I) DNA degradation by RhsB C-terminus (RhsB^CT^) and its mutant. The qualities of purified proteins are shown by SDS-PAGE analysis (right). (J) Competition assay of wild type (WT), Δ*tssM*, Δ*rhsB* and *rhsB^KE-AA^* mutant against the *B. subtilis* PY79. (K) Alignment of the predicted internal protease sequences between RhsB and TseI. (L) Western blotting analysis of RhsB^KE-AA^ and its cleavage-defective mutant D1484A. For panel C, E, G and J, error bars indicate the mean +/− standard deviation of three biological replicates and statistical significance was calculated using One-way ANOVA test for each group, \**P* < 0.05, \*\**P* < 0.01, \*\*\**P* < 0.001, \*\*\*\**P* < 0.0001.

To test if the C-terminus of RhsB (RhsB^CT^) is toxic, we expressed it using an arabinose inducible vector in *E. coli* and compared cell survival in the presence of arabinose (induced) or glucose (repressed). Results show that RhsB^CT^ is highly toxic, reducing survival by 100-fold when induced (Figure 2B). Although RhsB^CT^ belongs to the PD-(D/E)XK superfamily (13), there are no obvious catalytic sites in 4qbn or RhsB^CT^. We constructed several point mutations around the PD site and tested their toxicity in *E. coli*. Two mutants, KD-AA and KE-AA, were nontoxic while the K1561A mutant exhibited stronger toxicity than wild type RhsB^CT^ (Figure 2B). To test whether RimB proteins confer protection, we constructed the Δ*rhsB-rimB1&2* mutant lacking the immunity genes and transformed it with an empty pBBR vector or vectors encoding one or both immunity proteins (Figure 2C). Competition analyses against wild type AC or the Δ*tssM* mutant revealed that survival was significantly increased when immunity genes were ectopically expressed in the Δ*rhsB-rimB1&2* mutant, suggesting both immunity proteins could confer protection. Using bacterial two-hybrid assays, we found that both RimB1 and RimB2 could bind to the non-toxic RhsB^KE-AA^ construct, while the Aave_0500 protein, encoded downstream of *rimB2*, did not interact with any of the proteins (Figure 2D). As control, Pal and TolB proteins showed positive interaction.

Next, we tested whether RhsB is delivered to *B. subtilis* using a competition assay. The prey *B. subtilis* was transformed with the empty vector pHT01 or vectors expressing one or both immunity proteins, separately. Results show that all *B. subtilis* strains expressing the immunity genes survived significantly better than the one expressing the vector alone when they were competed with wild type AC (Figure 2E). When the Δ*rhsB* mutant was used as the killer, survival of *B. subtilis* with vector alone was increased to a similar level to that of *B. subtilis* expressing immunity genes. These results indicate that RhsB is delivered by the T6SS to *B. subtilis* and its toxicity is neutralized by the RimB immunity proteins expressed in *B. subtilis*.

To determine how RhsB is secreted, we used pull-down analysis and found that RhsB could direct interact with the upstream encoded VgrG3 and conserved chaperone EagT2, suggesting RhsB secretion is mediated by the VgrG spike complex (Figure 2F). Deletion of *vgrG3* resulted in significantly reduced killing of *B. subtilis* to a similar level to the *rhsB* mutant (Figure 2G). In addition, Western blotting analysis reveals that Hcp secretion was comparable among wild type, the Δ*rhsB*, and the Δ*vgrG3* (Figure 2H), eliminating the possibility that the reduced killing of Δ*rhsB* and Δ*vgrG3* is due to impaired T6SS secretion. Collectively, these results indicate that secretion of RhsB is mediated by VgrG3.

To test if RhsB exhibits nuclease activity, we purified the C-terminal domain of RhsB and RhsB^KD-AA^ mutant with an N-terminal 6His-SUMO tag. Results show that DNA was degraded by DNase I and wild type RhsB^CT^ but not by the RhsB^KD-AA_CT^ mutant (Figure 2I). Chromosomal inactivation of RhsB also attenuated the killing of *B. subtilis* to the same level as deletion of *rhsB* did (Figure 2J). In addition, sequence alignment of RhsB with TseI, a self-cleavable T6SS effector in *Aeromonas dhakensis* (14), shows that RhsB also contains the conserved aspartic catalytic residues for C-terminal cleavage (Figure 2K). We thus mutated one of the catalytic residues D1484 to alanine in the non-toxic RhsB^KE-AA^ background. Western blotting analysis shows that the C-terminus of RhsB was also cleaved by the internal protease and the D1484A mutation abolished the cleavage (Figure 2L).

In conclusion, we report the potent activities of *A. citrulli* T6SS against a panel of bacterial and fungal species. These findings have several important implications. Firstly, in comparison with previous efforts focused on human pathogens, T6SS functions are substantially less understood in plant pathogens despite the vast number of effectors, distinct ecological environments, and enormous impact to world economy and food security. Our findings not only address a long-standing critical question by showing T6SS can kill gram-positive species but also likely stimulate further research in novel T6SS functions of plant pathogens. Secondly, both the composition and physical size of cell envelopes of *B. subtilis* and *M. smegmatis* are quite distinct from the known susceptible gram-negative prey cells (15). Since expressing immunity proteins in *B. subtilis* confers protection against the delivered RhsB nuclease, this provides the first direct evidence that the T6SS of AC could penetrate into the cytosol of gram-positive cells. Notably, other effectors provide redundant functions to RhsB in killing *B. subtilis*, as evidenced by the attenuated but not abolished killing activities of the Δ*rhsB*. Further research is required to examine this penetration process in detail to elucidate the cellular death pathways using combinatorial effector mutations. Lastly, the fact that *A. citrulli* T6SS can kill both gram-negative and gram-positive bacterial and fungal species highlights the potential of developing T6SS-based treatment strategies as green alternatives to chemical agents in mitigating infectious diseases in agricultural and medical applications. In short, there seems to be no barrier too thick for the T6SS to break in the microbial world.

## Materials and Methods

### Bacterial strains and Growth conditions

All strains were routinely grown in LB, 7H9, 7H10 or YPD media following standard culturing conditions for each species. Antibiotics were used at the following concentrations: kanamycin (25 μg/ml for bacterial strains, 200 μg/ml for *S. cerevisiae* and 50 μg/ml for all other fungal strains), chloramphenicol (10 μg/ml), irgasan (25 μg/ml), and gentamicin (20 μg/ml), ampicillin (50 μg/ml). Gene expression vectors were constructed as previously described. All constructs were verified by sequencing. All plasmids and primers are available upon request.

### Bacterial cell killing assay

One milliliter of exponential phase cultures of killer cells (OD_600_=2) was centrifuged at 10,000 × *g* for 3 min and the resultant pellet was resuspended in fresh LB. For AC intraspecies competition, cells of killer and prey were mixed at a ratio of 20:1 (killer: prey) and spotted on LB agar plates for 24 h at 28 °C. For the interspecies competition, cells of killer and prey were mixed at a ratio of 2:1 (killer: prey) for *M. smegmatis* or 10:1 (killer: prey) for all other strains and spotted on 7H10 plates for *M. smegmatis* or LB agar plates for all other strains and coincubation for 8 h (for *M. smegmatis*), 10 h (for *S. cerevisiae*), or 3 h for all other strains at 37 °C (for all the bacterial prey cells) or 28 °C (for all the fungus prey cells) unless stated otherwise. Survival of prey cells was quantified by serial dilution and plating on selective media. Error bars show mean +/− standard deviation of three biological replicates.

### Western blotting analysis

Proteins were resolved on an SDS-PAGE gel and transferred to a PVDF membrane (Bio-Rad) by electrophoresis. The membrane was blocked with 5% [w/v] non-fat milk in Tris-buffered saline with TBST buffer (50 mM Tris, 150 mM NaCl, 0.1% [v/v] Tween-20, pH 7.6) for 1 h at room temperature, incubated sequentially with primary antibodies and secondary HRP-conjugated antibodies in TBST with 1% [w/v] milk. Signals were detected using the Clarity ECL solution (Bio-Rad). Monoclonal antibodies to epitope tags were from Thermo Scientific (V5, Product # 37-7500), and Biolegend (RpoB, Product # 663905). The polyclonal antibody to Hcp was custom-made by Shanghai Youlong Biotech. The HRP-linked secondary antibodies were purchased from ZSGB-Bio (Product # ZB-2305 (mouse) and # ZB-2301 (rabbit)).

### Protein secretion assay

Cultures were grown aerobically in LB with appropriate antibiotics at 37 °C to OD_600_ ~ 2 and collected by centrifugation at 2,500 × *g* for 8 min. Pellets were resuspended in fresh LB and incubated at 37 °C for 1 h. Cells were centrifuged at 10,000 × *g* for 2 min twice at room temperature. Pellets were resuspended in SDS-loading dye and used as whole-cell samples. Supernatants were precipitated in 20% [v/v] TCA (trichloroacetic acid) at −20 °C for 30 min. Samples were centrifuged at 15,000 × *g* for 30 min at 4 °C and pellets were washed with acetone, air-dried, and resuspended in SDS-loading dye. Whole-cell and secretion samples were boiled for 10 min for SDS-PAGE and Western blotting analysis. For secretome analysis, the supernatants of overnight cultures were collected and subject to SDS-PAGE analysis. Gel slices containing secreted proteins were cut out and sent for LC-MS/MS analysis performed by the Instrumental Analysis Center of Shanghai Jiao Tong University.

### Protein purification and enzymatic assays

Genes of interest were cloned into the pETSUMO vectors and transformed to *E. coli* BL21(DE3). Cells were grown in LB with appropriate antibiotics to OD_600_ ~ 0.6 at 37 Δ. Protein expression was induced with 1 mM IPTG at 37 ◻ for 5 h. The cells were centrifuged at 4,500 × *g* for 10 min. The pellets were resuspended in lysis buffer (20 mM Tris-HCl pH 8.0, 150 mM NaCl, 10 mM imidazole) and lysed by sonication. Lysates were centrifuged at 15,000 × *g* for 20 min and the supernatants were transferred onto Ni-NTA resin (Smart-lifesciences). Proteins were eluted in elution buffer (20 mM Tris-HCl pH 8.0, 150 mM NaCl, and variable concentrations of imidazole). Eluted samples were analyzed by SDS–PAGE.

Protein activity *in vitro* was detected by incubating with 30 ng plasmid at 37°C for 15 min. NEB CutSmart buffer (50 mM Potassium Acetate, 20 mM Tris-acetate, 10 mM Magnesium Acetate, 100 μg/ml BSA, pH 7.9) was chosen as the reaction buffer. Purified (0.1 μg) and 0.5 units DNase ◻ (positive control) were used separately in each reaction.

### Protein pull-down assays

Genes were cloned with epitope tags into pET and pBBRT vectors for expression. Cells were grown in LB with appropriate antibiotics to exponential phase, and induced with 1 mM IPTG overnight at 20 ◻ for pET vectors and 100 ng/ml aTc for 3 h at 37 ◻. Cells were collected by centrifugation, resuspended in lysis buffer (20 mM Tris pH 8.0, 500 mM NaCl, 50 mM imidazole with protease inhibitor (Thermo Scientific)), and lysed by sonication. After removing cell debris by centrifugation, supernatants were mixed and loaded to Ni-NTA resin (Smart-lifesciences), washed five times with wash buffer (20 mM Tris pH 8.0, 500 mM NaCl, 50 mM imidazole), and eluted in elution buffer (20 mM Tris pH 8.0, 500 mM NaCl, 500 mM imidazole). Input and eluted samples were analyzed by Western blotting.

### Protein toxicity assay

Cells expressing different plasmid constructs were grown in LB at 30 °C overnight supplemented with 0.2% [w/v] glucose. Cells were then collected and resuspended in fresh LB and grown to OD_600_ = 1. A series of 10-fold dilutions were plated on LB plates containing 0.1% [w/v] L-arabinose or 0.2% [w/v] glucose as indicated. Each experiment was repeated three times, with one representative experiment shown.

### Bacterial two-hybrid assay

Proteins of interest were fused to the T18 and T25 split domains of the *Bordetella* adenylate cyclase. The two plasmids encoding the fusion proteins were co-transformed into the reporter strain BTH101. Three independent colonies for each transformation were inoculated into 300 μl of LB medium. After 4 h growth at 30 °C, 3 μl of each culture were spotted onto LB plates supplemented with ampicillin, kanamycin, IPTG (0.02 mM), and X-Gal (40 μg/ml) and incubated for 6 h at 30 °C and then 10 h at room temperature. The experiments were done in triplicate and a representative result is shown.

### Bioinformatic analysis

All gene sequences of *A. citrulli* AAC00-1 are retrieved from the draft genome assembly (GenBank NC_008752.1). Benchling was used to manage and analyze DNA. RhsB sequence was analyzed with Phyre2. Homologs were aligned using Clustal Omega.

## Acknowledgments

This work was supported by funding from National Key R&D Program of China (2020YFA0907200), National Natural Science Foundation of China (31770082 and 32030001), Canadian Institutes of Health Research, Natural Sciences and Engineering Research Council of Canada, and Canada Research Chair program. We thank Steve Hersch and Kevin Manera for proofreading and helpful discussions. The funders had no role in study design, data collection and interpretation, or the decision to submit the work for publication.

## Author contributions

T.D. conceived the project. T.P., Y.K., Z.W., S.Y., M.T., H.L., Y.C., H.L., and H.Z. performed research; T.D. wrote the manuscript with assistance from T.P.

Correspondence and request for materials should be addressed to T. Dong. The authors declare no competing interests.

## References

1. S. T. Cole, et al., Deciphering the biology of *Mycobacterium tuberculosis* from the complete genome sequence. Nature 393, 537–544 (1998).

2. C. J. Nobile, A. D. Johnson, *Candida albicans* biofilms and human disease. Annu. Rev. Microbiol. 69, 71–92 (2015).

3. S. Pukatzki, et al., Identification of a conserved bacterial protein secretion system in *Vibrio cholerae* using the *Dictyostelium* host model system. Proc. Natl. Acad. Sci. 103, 1528–1533 (2006).

4. B. T. Ho, T. G. Dong, J. J. Mekalanos, A view to a kill: the bacterialtype VI secretion system. Cell Host Microbe 15, 9–21 (2014).

5. J. Wang, M. Brodmann, M. Basler, Assembly and subcellular localization of bacterial type VI secretion systems. Annu. Rev. Microbiol. 73, 621–638 (2019).

6. K. Trunk, et al., The type VI secretion system deploys antifungal effectors against microbial competitors. Nat. Microbiol. 3, 920–931 (2018).

7. A. T. Ma, S. McAuley, S. Pukatzki, J. J. Mekalanos, Translocation of a *Vibrio cholerae* type VI secretion effector requires bacterial endocytosis by host Cells. Cell Host Microbe 5, 234–243 (2009).

8. R. D. Hood, et al., A type VI secretion system of *Pseudomonas aeruginosa* targets a toxin to bacteria. Cell Host Microbe 7, 25–37 (2010).

9. S. Burdman, R. Walcott, *Acidovorax citrulli*: Generating basic and applied knowledge to tackle a global threat to the cucurbit industry. Mol. Plant Pathol. 13, 805–815 (2012).

10. A. Hachani, L. P. Allsopp, Y. Oduko, A. Filloux, The VgrG proteins are “à la Carte” delivery systems for bacterial type VI effectors. J. Biol. Chem. 289, 17872–17884 (2014).

11. T. G. Dong, B. T. Ho, D. R. Yoder-Himes, J. J. Mekalanos, Identification of T6SS-dependent effector and immunity proteins by Tn-seq in *Vibrio cholerae*. Proc. Natl. Acad. Sci. 110, 2623–2628 (2013).

12. S. Pennell, et al., FAN1 activity on asymmetric repair intermediates is mediated by an atypical monomeric virus-type replication-repair nuclease domain. Cell Rep. 8, 84–93 (2014).

13. K. Steczkiewicz, A. Muszewska, L. Knizewski, L. Rychlewski, K. Ginalski, SURVEY AND SUMMARY: Sequence, structure and functional diversity of PD-(D/E)XK phosphodiesterase superfamily. Nucleic Acids Res. 40, 7016–7045 (2012).

14. T.-T. Pei, et al., Intramolecular chaperone-mediated secretion of an Rhs effector toxin by a type VI secretion system. Nat. Commun. 11, 1865 (2020).

15. T. J. Silhavy, D. Kahne, S. Walker, The bacterial cell envelope. Cold Spring Harb. Perspect. Biol. 2, a000414 (2010).

